# Complete *de novo* assembly of *Tricholoma bakamatsutake* chromosomes revealed the structural divergence and differentiation of *Tricholoma* genomes

**DOI:** 10.1101/2023.02.12.528224

**Authors:** Hiroyuki Ichida, Hitoshi Murata, Shin Hatakeyama, Akiyoshi Yamada, Akira Ohta

**Author notes:** Corresponding author Hiroyuki Ichida, RIKEN Nishina Center for Accelerator-Based Science, 2-1 Hirosawa, Wako, Saitama 351-0198, Japan.

## Abstract

*Tricholoma bakamatsutake*, which is an edible ectomycorrhizal fungus associated with Fagaceae trees, may have diverged before the other species in *Tricholoma* section *Caligata*. We generated a highly contiguous whole-genome sequence for *T. bakamatsutake* SF-Tf05 isolated in an oak (*Quercus salicina*) forest in Japan. The assembly of high-fidelity long reads, with a median read length of 12.3 kb, resulted in 13 chromosome-sized contigs comprising 142,068,211 bases with an average GC content of 43.94%. The 13 chromosomes were predicted to encode 11,060 genes. A contig (122,566 bases) presumably containing the whole circular mitochondrial genome was also recovered. The chromosome-wide comparison of *T. bakamatsutake* and *T. matsutake* (TMA_r1.0) indicated that the basic number of chromosomes (13) was conserved, but the structures of the corresponding chromosomes diverged, with multiple inversions and translocations. Gene conservation and cluster analyses revealed at least three groups in *Tricholoma*. Specifically, all *T. bakamatsutake* strains belonged to the “bakamatsutake” clade, which is most proximal to the “caligatum” clade consisting of *T. caligatum* and *T. fulvocastaneum*. The constructed highly contiguous telomere-to-telomere genome sequence of a *T. bakamatsutake* isolate will serve as a fundamental resource for future research on the evolution and differentiation of *Tricholoma* species.

## Introduction

*Tricholoma* is a monophyletic genus comprising ectomycorrhizal fungi within the family Tricholomataceae (Moncalvo *et al*., 2002). *Tricholoma* species are distributed worldwide, but are mostly found in the temperate and subtropical zones of the southern and northern hemispheres. All known *Tricholoma* species are confirmed or predicted to be ectomycorrhizal fungi that are mainly associated with trees in the families Pinaceae, Betulaceae, and Fagaceae (Christensen & Heilmann-Clausen, 2013; Heilmann-Clausen *et al*., 2017). *Tricholoma matsutake* and its allied species *Tricholoma bakamatsutake* are valuable edible mushrooms that grow in Pinaceae and Fagaceae forests, respectively. Because these fungi, which belong to *Tricholoma* section *Caligata*, have limited micromorphological variations, they were historically often classified in the wrong taxa, but they were later reclassified according to molecular analyses (Trudell *et al*., 2017; Aoki *et al*., 2022).

*Tricholoma bakamatsutake* is a Fagaceae-associated mycorrhizal symbiont that often forms a mycelial colony known as ‘*shiro*’ in the A_0_ soil layer containing litter in both deciduous and evergreen broad-leaved forests, including those with trees from the genera *Quercus, Castanopsis* and *Pasania*, on which fruiting occurs (Ogawa, 1978; Ogawa & Ohara, 1978; Terashima, 1993; Terashima *et al*., 1993; Herrera *et al*., 2022). Unlike *T. bakamatsutake, T. matsutake*, which is a mycorrhizal symbiont of Pinaceae trees, such as those belonging to the genera *Pinus, Tsuga, Picea*, and *Abies*, colonizes the B soil layer mainly composed of minerals and rocks, with relatively little organic compounds (Yamada *et al*., 2006; Yamada *et al*., 2014; Endo *et al*., 2015). In addition to this ecological difference, *T. bakamatsutake* has a genetic trait that distinguishes it from *T. matsutake*. More specifically, it frequently generates phenotypic variants on agar plates, changing from a slow-growing brown mycelium to a fast-growing white mycelium. The former produces many chlamydospores with thick hyphae, whereas the latter produces relatively few chlamydospores with thin hyphae (Murata *et al*., 2022). On the basis of genomic analyses of other agaricomycetes, including the model ectomycorrhizal symbiont *Laccaria bicolor*, ectomycorrhizal fungi may have evolved later than saprophytic fungi (Labbé *et al*., 2012). There has been considerable interest in the evolution and differentiation of the genus *Tricholoma* in terms of the adaptations to different environments as well as the industrial utility of uncultivated prized mushrooms.

Retrotransposons are one of the most abundant repeating elements in the genomes of eukaryotic organisms, including plant-associated filamentous fungi (Dobinson & Hamer, 1993; Spanu *et al*., 2010; Labbé *et al*., 2012). Under the influence of environmental stimuli, retrotransposons replicate through RNA-mediated mechanisms and their DNA copies are integrated into the genome, leaving evolutionary footprints on the genome. In *Tricholoma*, three types of full-length retrotransposons have been identified, namely *marY1, marY2N*, and *megB1*, of which, the first two are exclusive to this genus. In addition, *marY1* is approximately 6 kb long and includes a 426 bp long terminal repeat (LTR) designated as *σ*_*marY1*_, which structurally resembles mammalian retroviruses. Moreover, it is transcribed and undergoes post-replication transposition (Murata & Yamada, 2000; Murata & Miyazaki, 2001; Murata & Miyazaki, 2004). In contrast, *marY2N* is a long interspersed nuclear element (LINE) that structurally resembles mRNA with a poly-(A) tail (Murata *et al*., 2001) and *megB1* is a dimeric *Alu*-like element (e.g., *AbaMEG1–AbaMEG2*). Earlier research showed that *Alu* is a short mobile DNA sequence with no coding region to replicate and integrate and is abundant in the form of a dimer in humans and primates (Deininger, 1989; Kazazian, 2000). Many agaricomycete fungi carry *megB1* in rRNA gene intergenic spacer 1 (IGS1) as the main component and multicopy element, but the fungi belonging to *Tricholoma* section *Caligata* carry a copy of *megB1* outside of the rRNA gene (Babasaki *et al*., 2007). On the basis of the copy numbers of these retrotransposons and the short repeating sequences surrounding *magB1*, the diversification of the species in *Tricholoma* section *Caligata* may have occurred as follows: *T. bakamatsutake* differentiated first, followed by Fagaceae-associated *T. fulvocastaneum* and then *T. matsutake* (Murata *et al*., 2013b).

The genome structures and sequences of most *Tricholoma* species have not been elucidated, but there have been several recent attempts at determining the *T. matsutake* whole-genome sequence (Min *et al*., 2020; Miyauchi *et al*., 2020). A telomere-to-telomere *T. matsutake* genome sequence was reported, but the sequencing was performed using two fruiting bodies collected under natural conditions and the sequenced fungus is not available as a strain (Kurokochi *et al*., 2022). There is little available genomic information regarding the phenotypically unstable *T. bakamatsutake*, which has ecological traits distinct from those of *T. matsutake* and may have diverged earlier than many other species belonging to the *Tricholoma* section *Caligata*. In the present study, we performed a whole-genome sequencing analysis of *T. bakamatsutake* strain SF-Tf05, which was originally collected in Shiga, Japan and has been stably maintained on agar for more than 10 years without any observable morphological changes. We constructed a telomere-to-telomere assembly of the strain using highly accurate long-read HiFi sequencing technology (Pacific Biosciences, Menlo Park, CA, USA). The resulting assembly consisted of 13 megabase-sized chromosomal contigs with a total length of 148.26 Mb. The number of chromosomes (13), overall GC content (43.94%), and high repetitive sequence content (78.68%) are consistent with the corresponding data for the previously reported *T. matsutake* assembly TMA_r1.0. However, the structure and/or sequence of the centromeric regions differ. Instead of general LINE-type sequences (e.g., those in *T. matsutake*), *marY2N* and its derivatives are enriched in the probable centromeric regions. The high-quality *T. bakamatsutake* SF-Tf05 genome sequence will serve as the foundation for future investigations on the evolution and differentiation of *Tricholoma* section *Caligata* and its related taxa.

## Materials and methods

### Fungal strains and growth conditions

The fungal strains used in this study are listed in Table 1. Unless otherwise stated, mycelia were cultured at 23 °C on modified Melin–Norkrans (MMN) medium containing 1.5% V8 juice (MMN+V8; Murata *et al*., 1999).

**Table 1.**
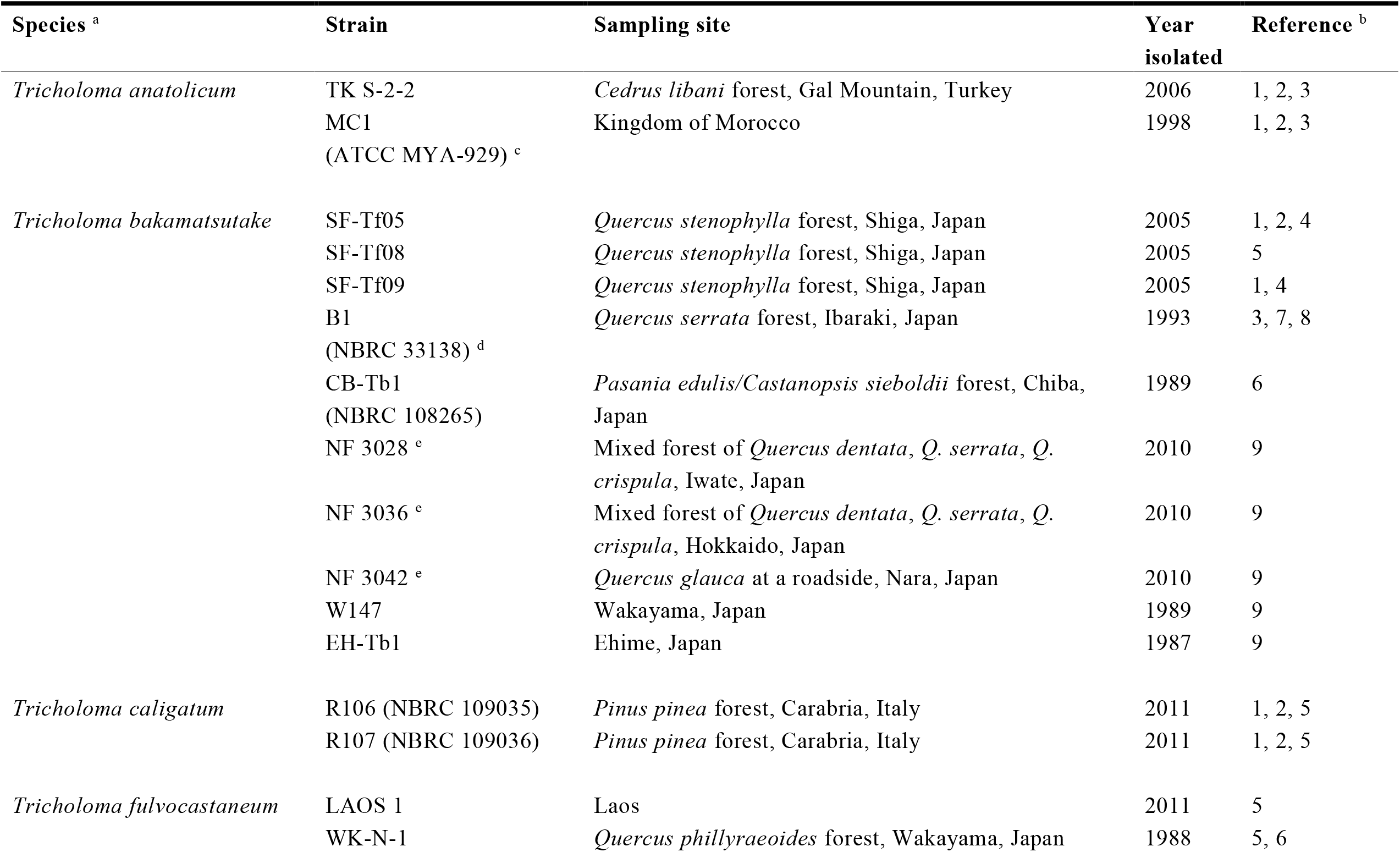
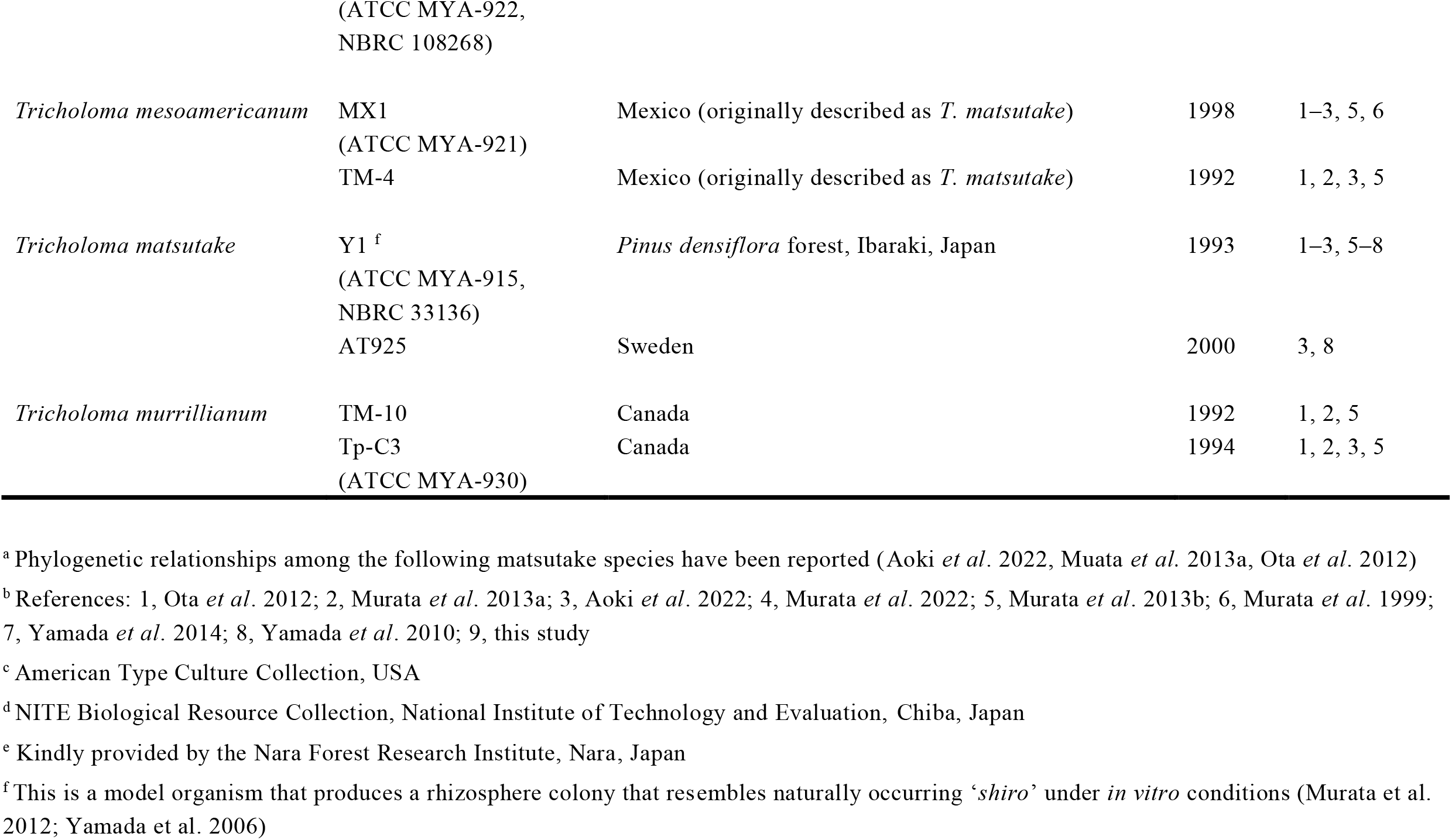
Fungal strains used in this study

### DNA extraction and sequencing

Genomic DNA was extracted from frozen mycelia using Genomic-tip (Qiagen, Hilden, Germany), with minor modifications to the manufacturer’s protocol. Briefly, 500 mg frozen mycelial mat was ground in liquid nitrogen using a mortar and pestle. The resulting fine powder was resuspended in 5 mL Buffer G2. After adding 10 µL RNase A (100 mg/mL, Qiagen) and 250 µL Proteinase K (20 mg/mL, Wako Pure Chemical, Osaka, Japan), the sample was incubated for 30 min at 55 °C with gentle agitation. Next, 3 mL chloroform was added and the solution was gently mixed in a rotator for 10 min. After centrifuging at 10,000 × g for 10 min at 4 °C, the supernatant was applied to a Genomic-tip 100/G and processed according to the manufacturer’s protocol. The resulting DNA was resuspended in 1× IDTE buffer (Integrated DNA Technologies, Coralville, IA, USA) and quantified using the QuantiFluor One dsDNA kit (Promega, Madison, WI, USA). The HiFi reads were obtained using the SMRT Cell 8M and the PacBio Sequel II instrument, which was operated in the circular consensus sequencing mode (Pacific Biosciences). The library was constructed and sequenced at the Genomics and Cell Characterization Core Facility (GC3F) at the University of Oregon (Eugene, OR, USA).

### Genome assembly and bioinformatics analysis

The default parameters of all programs were used unless otherwise specified. The HiFi reads were assembled using hifiasm (version 0.16.1-r375; Cheng *et al*., 2021). Smaller contigs (i.e., not including the 13 largest contigs) were subjected to a homology search, with BLASTN (version 2.12.0+; Altschul *et al*., 1997) used to screen the *Tricholoma* sequences in GenBank to identify possibly misassembled sequences and mitochondrial contigs. A Benchmarking Universal Single-Copy Orthologs (BUSCO; Simao *et al*., 2015) analysis using the Docker container (version 5.4.3_cv1) was performed to evaluate the completeness of the genome assembly. Telomere regions were identified by a BLASTN search involving five tandem repeats of the reported fungal 6-mer telomeric repeating unit (5′-CCCTAA-3′; Červenák *et al*., 2020; Rahnama *et al*., 2021), with an E-value cutoff of 1.0e^−10^. The *nucmer* program in the MUMmer4 suite (Marcais *et al*., 2018) was used for comparing genomes. The matching regions were filtered using the “-l 10000” option of the *delta-filter* program and plotted using Gnuplot (version 5.4 patchlevel 2).

Chromosomal genes were predicted using the FunGAP pipeline (version 1.1.1; Min *et al*., 2017) with RNA-seq reads from *T. matsutake* 945 deposited in the Sequence Read Archive (accession code PRJNA200596). The predicted genes were annotated according to a homology search of the NCBI non-redundant protein (nr) database (downloaded from https://ftp.ncbi.nlm.nih.gov/blast/db/FASTA) using the following options of the DIAMOND program (Buchfink *et al*., 2021): “--evalue 1e-10 --more-sensitive --query-cover 50 --subject-cover 50”. To functionally annotate genes, the KEGG Automatic Annotation Server (KAAS; https://www.genome.jp/kegg/kaas) was used along with an SBH method, GHOSTZ for the homology search, and the GENES dataset for 40 species (hsa, mmu, rno, dre, dme, cel, ath, sce, ago, cal, spo, ecu, pfa, cho, ehi, eco, nme, hpy, bsu, lla, mge, mtu, syn, aae, mja, ape, fox, mgr, ncr, bfu, ani, aor, ang, tms, ppl, tvs, hir, mpr, scm, and uma). The mitochondrial genome was annotated using MITOS2 (Donath *et al*., 2019), with “RefSeq 89 Fungi” as the reference and the “Mold” genetic codes.

### Repeated sequence and synteny analyses

Transposable elements and other repeated sequences were identified *ab initio* using RepeatModeler2 (version 2.0.3; Flynn *et al*., 2020), with the “-LTRStruct” option. Repetitive sequences within the chromosome assembly were identified using RepeatMasker (version 4.1.2-p1; developed by Smit, A.F.A., Hubley, R., and Green, P., downloaded from https://www.repeatmasker.org/), with the “-xsmall” option. Repeats were classified on the basis of the RepeatMasker output. The genomic distributions of *marY1* (GenBank accession code AB028236) and *marY2N* (AB047280) were determined using RepeatMasker and FASTA-formatted sequences as the repeat libraries. The LINE, *marY2N*, and LTR contents were calculated as the percentage of the masked regions in 5 kb (LINE and *marY2N*) or 50 kb (LTR and *marY1*) windows and plotted using Gnuplot. The GC content of genomic regions was calculated using GCcalc.py (version 85b6ab5; downloaded from https://github.com/WenchaoLin/GCcalc), with window and step sizes of 50 kb, and plotted using Gnuplot. The resulting plots were combined into a figure using Adobe Illustrator CS6 (Adobe Inc., San Jose, CA, USA). Genes within non-repetitive regions were extracted using the ‘intersect’ tool in BedTools (Quinlan & Hall, 2010) and the Tbkm_v1 repeat library created with RepeatModeler. The corresponding gene regions between Tbkm_v1 and TMA_r1.0 were determined using TBLASTN, with an E-value cutoff of 1.0e^−10^. Hits with the highest bit score and lowest E-value were considered the corresponding regions between these assemblies and visualized using the ‘Advanced Circos’ tool in the TBtools program (Chen *et al*., 2020).

### Gene conservation and mapping analyses of Tbkm_v1 and TMA_r1.0

The gene regions predicted by the FunGAP program were used as the input for the gene conservation analysis. Short read sequences from different *Tricholoma* species were mapped to Tbkm_v1 and TMA_r1.0 as previously described. The resulting BAM files were used for the gene coverage analysis. Mapping rates were calculated using the ‘idxstat’ tool in SamTools (Danecek *et al*., 2021). We used a combination of in-house C++ programs to calculate the gene coverage in the two reference sequences for each sample. First, read depth matrices at each chromosomal position were created using the ‘*create_read_count_matrix*’ program and then the percentage of gene regions that were covered with three or more reads was calculated using the ‘*gff_coverage*’ program. The resulting gene coverage information for Tbkm_v1 and TMA_r1.0 was combined and subjected to a cluster analysis using the ‘heatmap’ function in the R statistical environment (R Core Team, 2002).

## Results

### Telomere-to-telomere assembly of *T. bakamatsutake* SF-Tf05 chromosomes

We used genomic DNA extracted from *T. bakamatsutake* SF-Tf05 mycelia grown in a liquid medium. The extracted DNA contained a detectable amount of partially degraded fragments, presumably because of the relatively long culture time required for this slow-growing strain (4–6 weeks for the exponential growth phase). A sequencing run with the SMRT Cell 8M and the PacBio Sequel II instrument yielded 30.24 Gb of highly accurate long reads (HiFi reads) with a minimum quality of Q_20_ (i.e., consensus base call accuracy of 99%) and a median read length (read N_50_) of 12,294 bp. The assembly statistics are provided in Table 2. The assembly of the genome using hifiasm (Cheng *et al*., 2021) resulted in a total of 201 contigs with a total length of 148.26 Mb and a contig N_50_ value of 12.43 Mb. The longest contig was 14.07 Mb.

**Table 2.** Statistics for the Tbkm_v1 assembly

| **Assembly** | **All contigs** | **Tbk<sub>m</sub>_v1** | **Mitochondrial genome (mt)** |
| --- | --- | --- | --- |
| # contigs | 201 | 13 | 1 |
| Total length (bp) | 148,260,793 | 142,068,211 | 122,566 |
| Largest contig (bp) | 14,073,764 | 14,073,764 | n.a. |
| GC content (%) | 43.90 | 43.94 | 21.70 |
| N50 | 12,427,164 | 12,427,164 | n.a. |
| N75 | 9,419,585 | 10,325,240 | n.a. |
| L50 | 6 | 6 | n.a. |
| L75 | 10 | 9 | n.a. |
| Ns per 100 kb | 0 | 0 | 0 |

n.a., not applicable

The 13 longest contigs represented 95.82% (142.07 Mb) of the total contig length. The lengths of the 13th and 14th contigs differed by more than 10-times (5.23 Mb and 0.31 Mb, respectively). Therefore, we considered the 13 longest contigs as the primary contigs and designated them as chromosomes 1–13 in descending order of their nucleotide length. The BLASTN search using the reported telomeric repeat sequence in fungi (5 ′-CCCTAA-3′; Červenák *et al*., 2020; Rahnama *et al*., 2021) revealed telomeric repeats on at least one side of all chromosomes. Nearly half of the *T. bakamatsutake* chromosomes (chromosomes 1, 4, 7, 9, 10, and 12) had telomeric repeats at both ends, indicating that most of the assembled sequences were telomere-to-telomere sequences. The assembled chromosome lengths and GC contents varied from 14.07 Mb to 5.23 Mb and from 43.39% to 44.52%, respectively (Table S1). The overall GC content of the 13 chromosomes was 43.94%, which was similar to the GC contents of previously reported *T. matsutake* genome assemblies, with an average of 45.39% in three different assemblies (Min *et al*., 2020; Miyauchi *et al*., 2020; Kurokochi *et al*., 2022). Unlike the *T. matsutake* chromosomes, most of the *T. bakamatsutake* chromosomal regions had similar GC contents. The exceptions were chromosomes 2 and 8, which had a GC-rich region near one of the ends. These observations suggested the physical structures of the centromeric regions may differ between these two species (Fig. 1). The BUSCO analysis indicated that the 13 chromosomes contained 97.7% (741 of 758) of the core genes defined in the fungi_odb10 dataset, suggesting the assembly is almost complete. The final chromosomal assembly was designated Tbkm_v1 and deposited in GenBank with the accession codes CP114857 –CP114869 (chr01–13).

**Figure 1.**
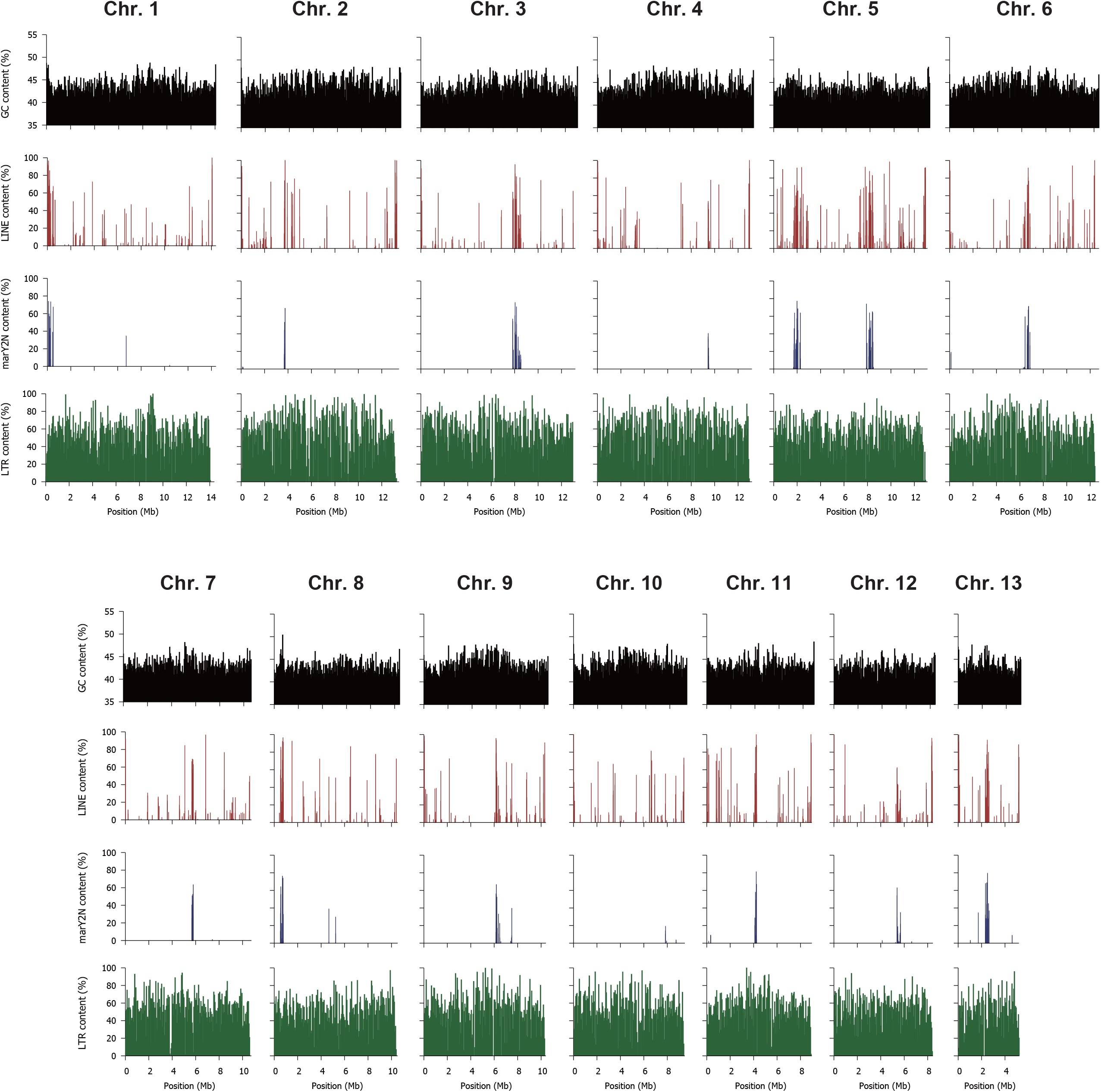
Characteristics of *T. bakamatsutake* SF-Tf05 chromosomes. The contents of guanine and cytosine nucleotides (GC), long interspersed nuclear elements (LINEs), *marY2N*, and long terminal repeat (LTR)-type retrotransposons for different chromosomes are presented. The window sizes are 50 kb for GC, LINE, and LTR and 5 kb for *marY2N*. Unlike in *T. matsutake*, GC- and LINE-rich regions were not detected in the chromosomal regions.

### Prediction of chromosomal genes

Protein-coding genes on chromosomes were predicted using the FunGAP2 pipeline (Min *et al*., 2017) and RNA-seq reads from two different analyses of *T. matsutake* (SRA BioProject codes PRJNA365663 and PRJNA536102). Although this was a heterologous mapping between different species, the mapping rate of the RNA-seq reads to the Tbkm_v1 assembly was 63.16% (a total of 314.07 million mapped reads), which was sufficient for creating gene models on the basis of the conserved genes. The FunGAP pipeline predicted 11,060 genes encoded by the 13 chromosomes, with an average gene density of one per 13.42 kb (range: one gene per 9.89–16.69 kb among chromosomes). The predicted proteins covered 96.7% (733 of 758) of the BUSCO groups defined in the fungi_odb10 dataset, reflecting the suitability of the method used for predicting genes. The predicted genes were annotated by using the deduced amino acid sequences as queries to screen the nr database for similar sequences, with a cutoff threshold of 50% coverage in both query and hit sequences. Of the 11,060 predicted gene products, 9,200 (83.18%) had a significant hit in the nr database. Approximately 80% (7,512 gene products; 83.18%) of these best hits in the nr database were predicted proteins in *T. matsutake* 945. Additionally, nearly 60% (5,416 gene products; 58.87%) were annotated as “hypothetical protein,” “uncharacterized protein,” and “unnamed protein product.” Although the remaining gene products also had unclear functions, they were mostly similar to cytochrome P450 (77 proteins), MFS general substrate transporters (74 proteins), and alpha/beta-hydrolases (58 proteins). The KAAS analysis mapped 4,802 predicted genes to the KEGG Orthology (KO) groups and assigned the genes to functional categories and biological pathways. Table S2 lists the number of genes in each functional group. In addition to the genes assigned to groups associated with housekeeping functions, such as membrane trafficking (ko04131; 411 genes) and messenger RNA biogenesis (ko03019, 204 genes), many genes were classified as transporters (ko02000, 209 genes), peptidases and inhibitors (ko01002, 149 genes), and protein kinases (ko01001, 119 genes). A total of 365 tRNA genes were identified by the tRNAscan-SE program, whereas another seven genes were classified as “tRNAs with undetermined/unknown isotypes.” Interestingly, Tbkm_v1 included more than the expected number of tRNA-Ile genes, most of which (275 of 277) harbored the TAT anticodon.

### Mitochondrial genome

The BLASTN searches using the remaining 188 contigs to screen the NCBI nucleotide database (nr/nt) revealed a contig (138,202 bp) that matched a previously reported complete mitochondrial genome sequence in *T. bakamatsutake* (GenBank accession code MN873035; 103,090 bp). The two mitochondrial genome sequences were highly similar (i.e., sequence identity exceeding 99%). The sequence comparison showed that our initially assembled mitochondrial sequence had overlapping sequences at each end, with the first 15,636 bp perfectly matching the sequence at the 3′ end. This overlap indicated that the mitochondrial genome was organized as a closed master circle. One of the two duplicated regions was manually removed. The remaining 122,566 bp sequence was designated as ‘ *mt*’ and considered to represent the complete *T. bakamatsutake* SF-Tf05 mitochondrial genome sequence.

Although the assembled *mt* sequence was highly similar to a previously reported mitochondrial genome sequence (MN873035), there were five major differences: an inversion between positions 9,623 and 22,404 (12,782 bp) as well as four insertions between positions 7,573 and 9,622 (2,050 bp), 22,405 and 31,465 (9,061 bp), 63,476 and 64,161 (686 bp), and 81,405 and 88,967 (7,563 bp). The Liftoff program (Shumate & Salzberg, 2020) mapped all of the 228 annotated features in MN873035, of which 213 were detected as single copies and the remaining 15 had an extra copy in the *mt* sequence. There were 81 protein-coding genes and 25 genes encoding tRNAs corresponding to 17 of the 20 amino acids in the SF-Tf05 mitochondrial genome. The sequence analysis indicated that the SF-Tf05 mitochondrial genome lacks tRNA genes for proline (AGG, GGG, CGG, and TGG), glutamic acid (CTC and TTC), and cysteine (ACA and GCA). The same tRNA gene analyses were also performed for MN873035 and AP026551, which are the complete mitochondrial genome sequences in *T. bakamatsutake* and *T. matsutake*, respectively. Similar to the results for *T. bakamatsutake* SF-Tf05, the MN873035 and AP026551 sequences did not include tRNA genes for proline, glutamic acid, and cysteine. In addition to the tRNAs for these three amino acids, MN873035 also lacked tRNAs for lysine, serine, and tyrosine. These observations revealed that some tRNA genes in *Tricholoma* species are encoded in the nuclear genome. To further investigate the completeness and possible sequencing and assembly errors in common mitochondrial genes, we performed a *de novo* gene prediction and annotation using MITOS2 (Donath *et al*., 2019) for both SF-Tf05 and MN873035. The results indicated the coding regions of the *dpo* and *lagli* genes in SF-Tf05 and MN873035 may include stop codons. Some additional stop codons were also detected in the SF-Tf05 *giy* gene. The Sanger sequencing of the PCR-amplified fragments for these regions in SF-Tf05 confirmed the presence of all of these mutations in the genome (data not shown). The resulting complete mitochondrial genome sequence was deposited in GenBank with the accession code CP114870.

### Repeated sequences in the *T. bakamatsutake* genome

The repeated sequences in the Tbkm_v1 assembly were identified using RepeatModeler and RepeatMasker. In the Tbkm_v1 assembly, 78.68% (111.78 Mb) of the entire genome consisted of repetitive sequences. The abundance of each repetitive sequence is summarized in Table 3. The LTR retrotransposons were the most abundant type of repeated sequences in Tbkm_v1 (i.e., 53.16% of the genome). Almost all of the LTR-type elements were classified as Ty3/Gypsy (74.67% of all LTR regions) or Ty1/Copia (19.55%). The other types of repeated sequences in Tbkm_v1 were mostly DNA transposons (6.66% of the genome), rolling circles (6.38%), and other unclassified interspersed repeats (10.82%). The total length of small RNA, simple repeats, and other low complexity regions represented less than 1% of the genome.

**Table 3.** Repetitive sequences in the Tbkm_v1 assembly

| **Type of repeated sequence** | **Num. of elements** | **Total length (bp)** | **Percentage of genome** |
| --- | --- | --- | --- |
| SINEs | 299 | 81,536 | 0.06 |
| LINEs | 1,736 | 1,756,689 | 1.24 |
| LTR elements | 58,477 | 75,528,114 | 53.16 |
| DNA Transposons | 10,103 | 9,464,515 | 6.66 |
| Rolling-circles | 4,969 | 9,030,370 | 6.36 |
| Unclassified interspersed repeats | 44,596 | 15,366,895 | 10.82 |
| Small RNA | 379 | 119,893 | 0.08 |
| Simple repeats | 9,742 | 473,500 | 0.33 |
| Low complexity | 674 | 40,320 | 0.03 |
| **Total** |  | **111,780,296** | **78.68** |

Compared with *T. matsutake*, the localization of LINEs was less extensive in Tbkm_v1, with multiple chromosomal regions that contained many LINEs. Moreover, the locations of LINE-rich regions were not highly correlated with the GC contents (Fig. 1). Some of these LINE-rich regions overlapped the localized regions containing *marY2N*-like sequences, which were previously characterized as LINE-like non-LTR retroelements. With the exception of chromosome 10, which did not have an apparent localized peak, the *marY2N*-rich regions were located only near the center or the end of chromosomes. These observations suggest that *marY2N* and its derivatives are enriched in centromeric regions. Another well-studied retroelement in *T. matsutake, marY1* (Murata & Yamada, 2000; Kurokochi *et al*., 2022), was distributed across all 13 chromosomes in Tbkm_v1 (Fig. S1). A BLASTN search using the full-length *marY1* sequence (GenBank accession code AB028236) as the query and an E-value cutoff of 1.0e^−10^ detected 559 copies of *marY1*-like sequences in the assembly.

### Mating type (*MAT*) loci in *T. bakamatsutake*

The mating type (*MAT*) genes have been extensively studied in many basidiomycetous fungi (Kües, 2015). The bipolar and tetrapolar systems are the two major types of systems that determine the mating type in Basidiomycota fungi. The two *MAT* loci, *MATA* (syn. *b* or *HD*) and *MATB* (syn. *a* or *P/R*), which comprise a cluster of homeodomain transcription factor genes and pheromone receptor (*STE3*) and precursor (*Phe3*) genes, are linked at a single *MAT* locus (bipolar system) or are separated, often on different chromosomes (tetrapolar system) (Peris *et al*., 2022). Homology searches using the deduced HD and STE3 protein sequences of *Hypsizygus marmoreus* (Wang *et al*., 2021) revealed that *T. bakamatsutake* harbors *HD* and *STE3* orthologs on different chromosomes, implying that the mating type in this species is determined by the tetrapolar system. The schematic structures of the *MATA* and *MATB* loci in *T. bakamatsutake* are presented in Fig. 2. The *MATA* locus (approximately 17.4 kb region on chromosome 2) is flanked by the mitochondrial intermediate peptidase (*Mip*) and beta-flanking (*Bfg*) genes, which is consistent with recent findings regarding *H. marmoreus* (Wang *et al*., 2021). Similar to *L. bicolor*, but unlike the two *Trichaptum* species with a sequenced genome and some other Basidiomycota fungi (James *et al*., 2004; Niculita-Hirzel *et al*., 2008; Peris *et al*., 2022), the glycogenin-1 (*GLGEN*) gene is located immediately next to *Bfg* in *T. bakamatsutake*. The region between *Mip* and *Bfg* was predicted to include four genes, of which the deduced amino acid sequence of *SF02g08880* matched HD2.1 of *H. marmoreus* (QQL12049), whereas *SF02g08870* and *SF02g08890* matched HD1.2 (QQL12048). The *MATB* locus (approximately 288.6 kb region on chromosome 11) contains six genes encoding STE3.s2-like proteins (*SF11g01200* and *SF11g01290–SF11g01330*) as well as individual genes encoding STE3.s1-, STE3.1-, and STE3.2-like proteins (*SF11g01280, SF11g01340*, and *SF11g01380*, respectively). An additional STE3-like gene (*SF11g01390*) was detected in *T. matsutake* (KAF8222657) and *Clitocybe nuda* (KAF9468417), an edible mushroom in the family Tricholomataceae, but not in *H. marmoreus*. The structures and gene organizations in the *MATA* and *MATB* loci were essentially conserved in *T. bakamatsutake* and *T. matsutake* (Fig. 2). These results indicated that mating compatibility is determined by a tetrapolar system in *T. bakamatsutake* and *T. matsutake*.

**Figure 2.**
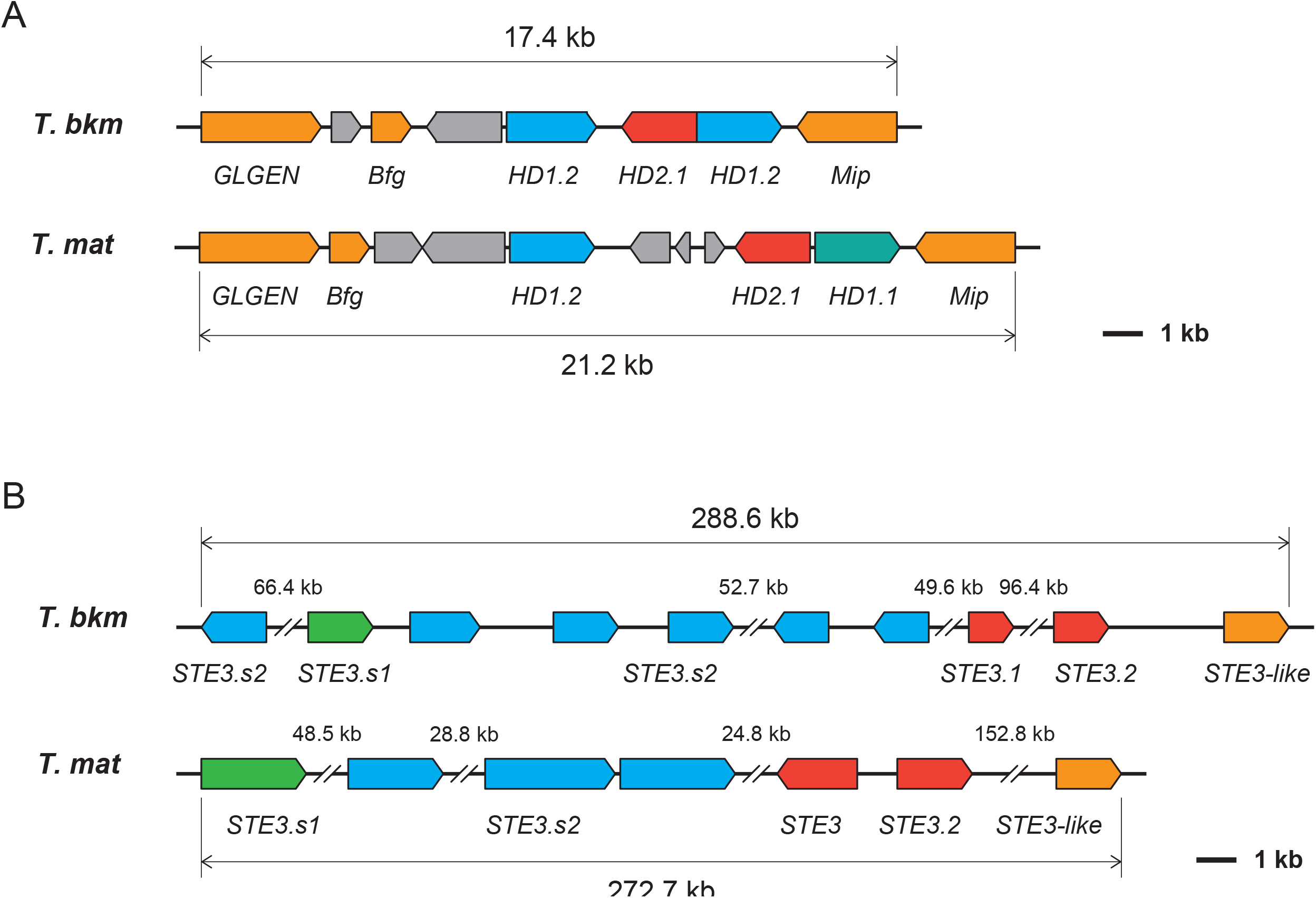
Structures of the mating type loci in *T. bakamatsutake* and *T. matsutake*. Gene organizations in the probable *MATA* (**A**) and *MATB* (**B**) loci in *T. bakamatsutake* (*T. bkm*) and *T. matsutake* (*T. mat*). Box colors reflect the similarity to known mating type genes. The genetic contents in the *MAT* regions are similar between the two *Tricholoma* species.

### Common and unique genes in the two *Tricholoma* species

We investigated the gene conservation between *T. bakamatsutake* and *T. matsutake* using the telomere-to-telomere assemblies Tbkm_v1 and TMA_r1.0. Although TMA_r1.0 was previously predicted to contain 28,322 protein-coding genes (Kurokochi *et al*., 2022), information regarding these genes was not publicly available when we conducted the present study.

Therefore, the genes in TMA_r1.0 were predicted and annotated according to the method used for Tbkm_v1. The FunGAP pipeline revealed 18,646 protein-coding genes encoded by the 13 chromosomes in TMA_r1.0. We subsequently used all of the predicted protein sequences from Tbkm_v1 and TMA_r1.0 to cross-search the matching region via a TBLASTN search with an E-value cutoff of 1.0e^−10^. There were 928 (8.39% of the predicted genes in Tbkm_v1) and 5,395 (28.94% of our predicted genes in TMA_r1.0) genes that were present in one of the two assemblies. Another 893 (8.07%) and 2,767 (14.84%) genes had deduced amino acid sequence similarities/identities less than 50% in Tbkm_v1 and TMA_r1.0, respectively. The remaining 9,239 (83.54%) and 10,484 (56.23%) genes were considered to be common to both species.

Even though more genes were predicted for TMA_r1.0 than for Tbkm_v1 because RNA-seq reads from the same species were used, it was surprising that nearly 30% of the genes were detected only in TMA_r1.0, while more than 80% of the genes in Tbkm_v1 were present in both species. The functional categories of the genes that were present in at least one of the species are presented in Fig. 3. Protein kinases (ko01001; 12 and 18 unique genes in Tbkm_v1 and TMA_r1.0, respectively), peptidases and their inhibitors (ko01002; 6 and 12 genes), transporters (ko02000; 6 and 31 genes), and amino acid metabolism (ko99985; 5 genes in Tbkm_v1) were the notable categories in addition to some of the large functional categories associated with the protein-coding genes in the genomes described above. Many of the species-specific genes were revealed to encode relatively small peptides and proteins: 216.23 ± 205.69 (average ± standard deviation) and 216.67 ± 194.14 in Tbkm_v1 and TMA_r1.0, respectively. These peptides and proteins were smaller than the common gene products in Tbkm_v1 and TMA_r1.0: 458.86 ± 367.83 and 437.81 ± 358.81, respectively.

**Figure 3.**
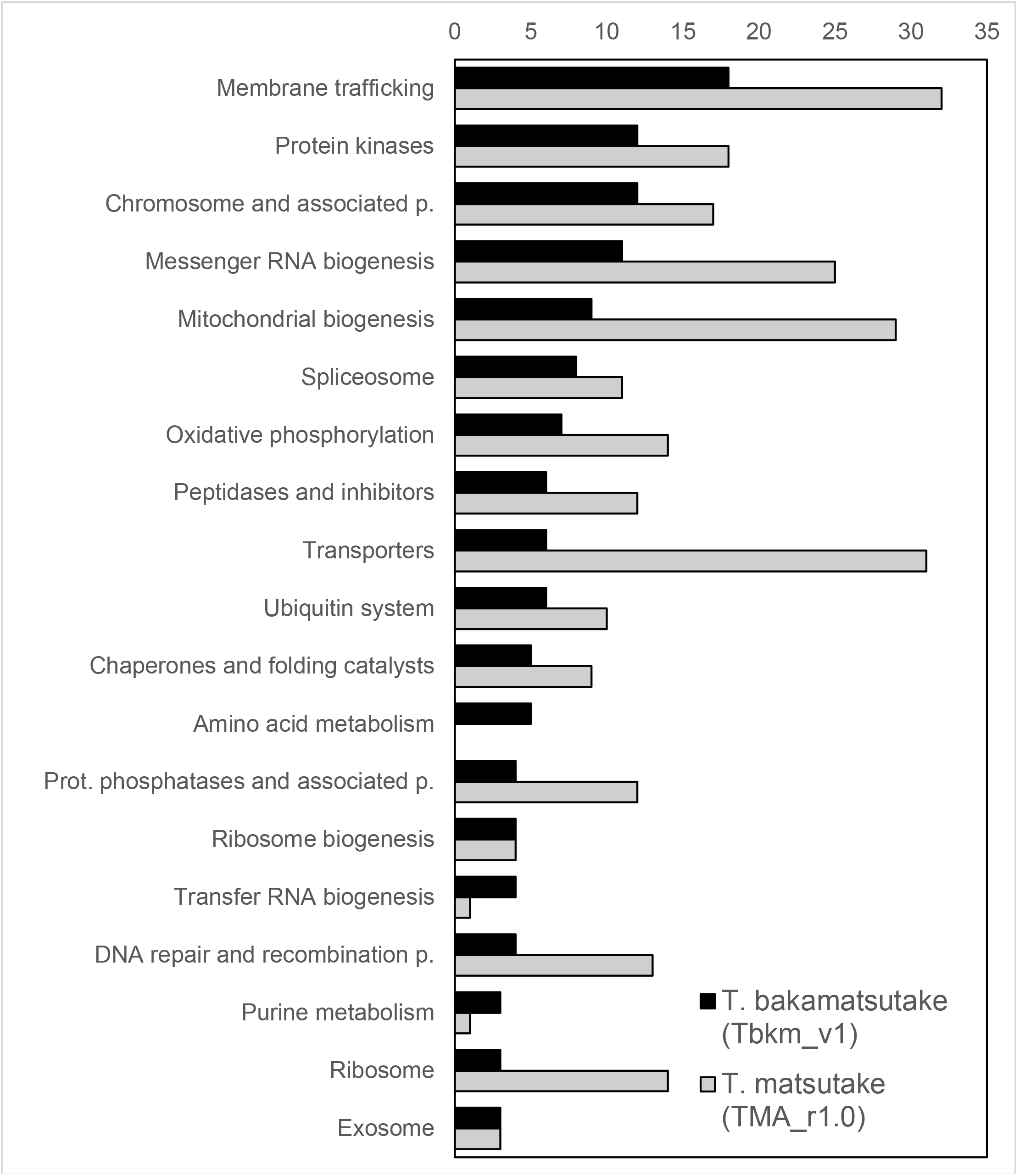
Functional categorization of genes that were present in either *T. bakamatsutake* (Tbkm_v1) or *T. matsutake* (TMA_r1.0). Bars indicate the number of genes in each functional category. Genes were categorized according to the KEGG Orthology database.

Multiple *T. bakamatsutake* genes were detected exclusively in Tbkm_v1. The functionally annotated genes included *SF01g07050* (encoding endo-1,4-β-xylanase), *SF12g05070* (encoding cholesterol 7-α-monooxygenase), *SF03g00020* and *SF04g00010* (encoding mitogen-activated kinase-like proteins), *SF01g08930, SF01g08940, SF01g09180, SF06g10950*, and *SF07g06230* (possibly encoding glutamine amidotransferase DUG3-like proteins according to the KAAS results), and five genes (*SF01g08930, SF01g08940, SF01g09180, SF06g10950*, and *SF07g06230*) that were significantly similar to the gene encoding the “hypothetical protein H0H93_006617” containing a YafJ-type glutamine amidotransferase-like domain (cd00352 in the NCBI Conserved Domain Database; Yang *et al*., 2020) from the basidiomycete *Arthromyces matolae*. In contrast, genes encoding AMY (α-amylase, *TM03g03410*), PHO (acid phosphatase, *TM01g00540*), and endo-β-glucanase-like (*TM05g15980*) were present only in TMA_r1.0. We determined that Tbkm_v1 lacks a gene encoding PTEN, which catalyzes the dephosphorylation of phosphatidylinositol 3,4,5-trisphosphate (PIP_3_) to produce phosphatidylinositol-4,5-bisphosphate (PIP_2_). Compared with other membrane phospholipids, PIP_2_ and PIP_3_ are less abundant and they function as second messengers in several crucial cellular processes, such as membrane trafficking and cell signaling, and are highly conserved across kingdoms. The phosphatase PTEN contributes to a negative regulatory mechanism that converts PIP_3_ back to PIP_2_, thereby decreasing the abundance of PIP_3_ in the plasma membrane (Czech, 2000). Theoretically, a lack of PTEN increases the PIP_3_ content and may lead to the continuous activation and deactivation of downstream processes. Therefore, it is possible that some of the presented genetic diversity was the result of the differences in the adaptations to environmental conditions between *T. bakamatsutake* and *T. matsutake*, which establish ectomycorrhizal symbiotic relationships with broad-leaved deciduous Fagaceae trees and coniferous Pinaceae trees, respectively.

### *T. bakamatsutake* has a unique nitrogen assimilation mechanism

On the basis of our comparisons, we determined that *T. bakamatsutake* SF-Tf05 lacks *nit-6* (KO: K17877), which encodes a nitrite reductase that catalyzes the conversion of nitrite to ammonia in an assimilatory nitrate reduction pathway. Another functional homolog (*nir1/nirA*) in this pathway encodes a ferredoxin-type nitrite reductase that is present mainly in plants and bacteria. However, an ortholog of *nir1/nirA* was not identified in Tbkm_v1 or TMA_r1.0.

Thus, *nit-6* may be the only nitrite reductase gene in *Tricholoma* species. Notably, *SF09g00300* in Tbkm_v1 encodes a protein similar to a nitrate/nitrite transporter identified in saprotrophic Agaricomycetes fungi, including *Cyathus striatus* (GenPept accession code KAF9007402) and *Mycena venus* (KAF7357751), as well as in many other species. In contrast, the *T. matsutake* genome, which contains *nit-6*, does not include an *SF09g00300* ortholog. Our PCR experiments demonstrated that the genomes of all 10 examined *T. bakamatsutake* isolates collected from different regions in Japan carry *SF09g00300*, but not *nit-6* (Fig. S2A). Conversely, the genomic analysis of 12 isolates of different *Tricholoma* species (i.e., *T. anatolicum, T. fulvocastaneum*,

*T. murrillianum*, and *T. matsutake*) revealed the presence of *nit-6* and the absence of *SF09g00300* (Fig. S2B and C). Interestingly, *T. caligatum* and *T. mesoamericanum* apparently lack both *nit-6* and *SF09g00300* (Fig. S2C). The analysis of the products of the PCR amplification using *nit-6* primers and *T. fulvocastaneum* strains LAOS1 (originally collected in Laos) and WK-N-1 (Japan) confirmed that the sequences did not match *nit-6*, indicative of non-specific PCR amplifications. We speculated that the probable nitrate/nitrite transporter encoded by *SF09g00300* mediates an alternative nitrogen assimilation mechanism in *T. bakamatsutake*. Additionally, the nitrogen assimilation mechanisms of *T. caligatum* and *T. mesoamericanum* may not involve *nit-6* and *SF09g00300*. These results suggest that *T. bakamatsutake* evolved a nitrite uptake and assimilation system that has not been observed in other *Tricholoma* species.

### Chromosome-scale synteny between Tbkm_v1 and TMA_r1.0

The telomere-to-telomere assemblies of all 13 chromosomes in *T. bakamatsutake* (Tbkm_v1) and *T. matsutake* (TMA_r1.0) enabled us to characterize the chromosome-wide syntenic regions between the two genomes. Because repetitive sequences occupied more than 70% of the complete genome in both Tbkm_v1 and TMA_r1.0, simple global alignments of chromosome or genome segments were not useful. Accordingly, we used the deduced amino acid sequences of the predicted genes in non-repetitive regions in Tbkm_v1 and performed TBLASTN searches to find the corresponding genomic regions in TMA_r1.0. Using this approach, multiple hits in different genomic loci due to dispersed repeated sequences were effectively eliminated. We identified and visualized the chromosome-wide synteny between these two species (Fig. 4 and Table S3). Chromosomes 1 and 13 in Tbkm_v1 (Tb1 and Tb13) had a chimeric structure that matched approximately half of chromosomes 10 and 12 in TMA_r1.0 (Tm10 and Tm12). There was an almost one-to-one correspondence among the other 11 chromosomes, but the corresponding chromosomes in the two assemblies were not sequential, even though the 13 chromosomes were numbered according to the descending order of their lengths in both assemblies. Moreover, the examination of the corresponding regions between Tb5/Tb11 and Tm1/Tm11 revealed multiple inversions within the chromosomes.

**Figure 4.**
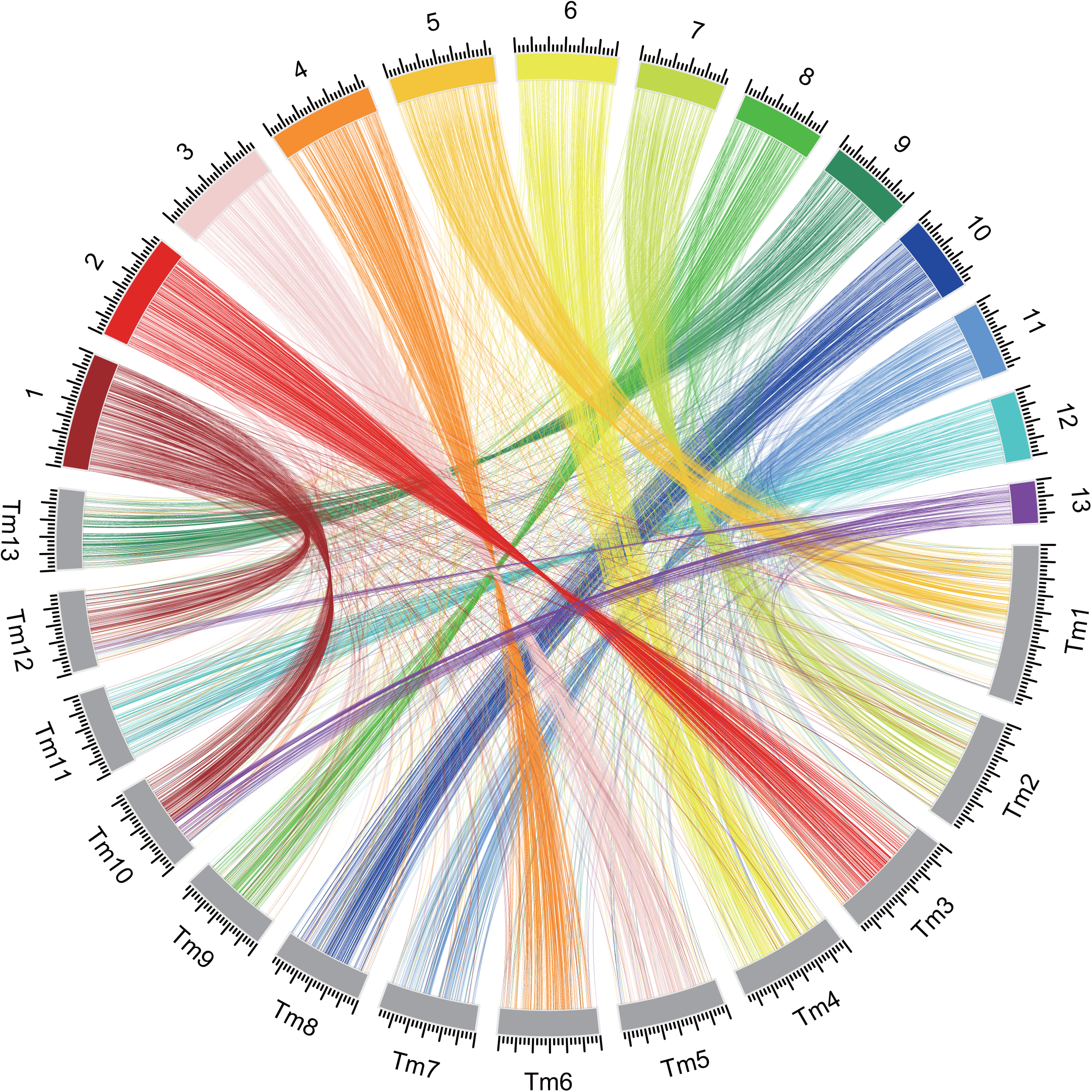
Chromosome-wide comparison of the *T. bakamatsutake* (Tbkm_v1) and *T. matsutake* (TMA_r1.0) genomes. TBLASTN searches were performed using the gene sequences located within non-repetitive regions as queries. The searches were restricted to matches with E-values of 1.0e^−10^ or less. The best-hit regions were considered as the corresponding regions. The results were visualized using the Circos software.

These results indicated that during species differentiation, *T. bakamatsutake* and *T. matsutake* independently accumulated different types of DNA in their chromosomes, while still maintaining the basic number of chromosomes. Such large alterations in chromosomal structures and multiple inversions within chromosomes may be related to the mechanisms underlying the reproductive isolation of these species.

### *Tricholoma* section *Caligata* consists of at least three major karyotypes

The comparison of the conserved genes in Tbkm_v1 and TMA_r1.0 detected the chromosome-wide synteny in these species. We then investigated the conservation of genes among *Tricholoma* section *Caligata* species. Briefly, we performed a random shotgun sequencing analysis of the following eight strains: *T. bakamatsutake* (SF-Tf05 and NBRC 33138), *T. matsutake* (NBRC 33138), *T. anatolicum* (MC1), *T. caligatum* (R107), *T. fulvocastaneum* (WK-N-1), *T. mesoamericanum* (MX1), and *T. murrillianum* (Tp-C3). The generated sequencing reads as well as one read for *T. matsutake* 945 obtained from SRA (BioProject code PRJNA200596) were mapped to the Tbkm_v1 and TMA_r1.0 sequences. The mapping rates are summarized in Table S4. Three different groups had high (>90%) and low (approximately 50%) fractions of mapped reads in the two assemblies. The first group consisted of the two *T. bakamatsutake* strains (high and low fractions of mapped reads in Tbkm_v1 and TMA_r1.0, respectively). The second group included *T. matsutake, T. anatolicum, T. mesoamericanum*, and *T. murrillianum* (high and low fractions of mapped reads in TMA_r1.0 and Tbkm_v1, respectively). The remaining two species, *T. caligatum* and *T. fulvocastaneum*, formed the third group (low fractions of mapped reads in Tbkm_v1 and TMA_r1.0). These findings implied that approximately half of the genomic regions were common to all of the examined *Tricholoma* species, but the other half differentiated into at least three groups. We calculated the read coverage (i.e., percentage of the gene regions that were covered by mapped reads) in both assemblies and used it as the input for the cluster analysis that used Euclidean distances as the distance matrix. The cluster analysis also revealed three major clades (Fig. 5). In the “matsutake” clade, *T. anatolicum* and *T. mesoamericanum* were most similar to *T. matsutake*, followed by *T. murrilianum*. The *T. bakamatsutake* strains formed their own “bakamatsutake” clade, which was proximal to the third clade “caligatum,” which consisted of *T. caligatum* and *T. fulvocastaneum*.

**Figure 5.**
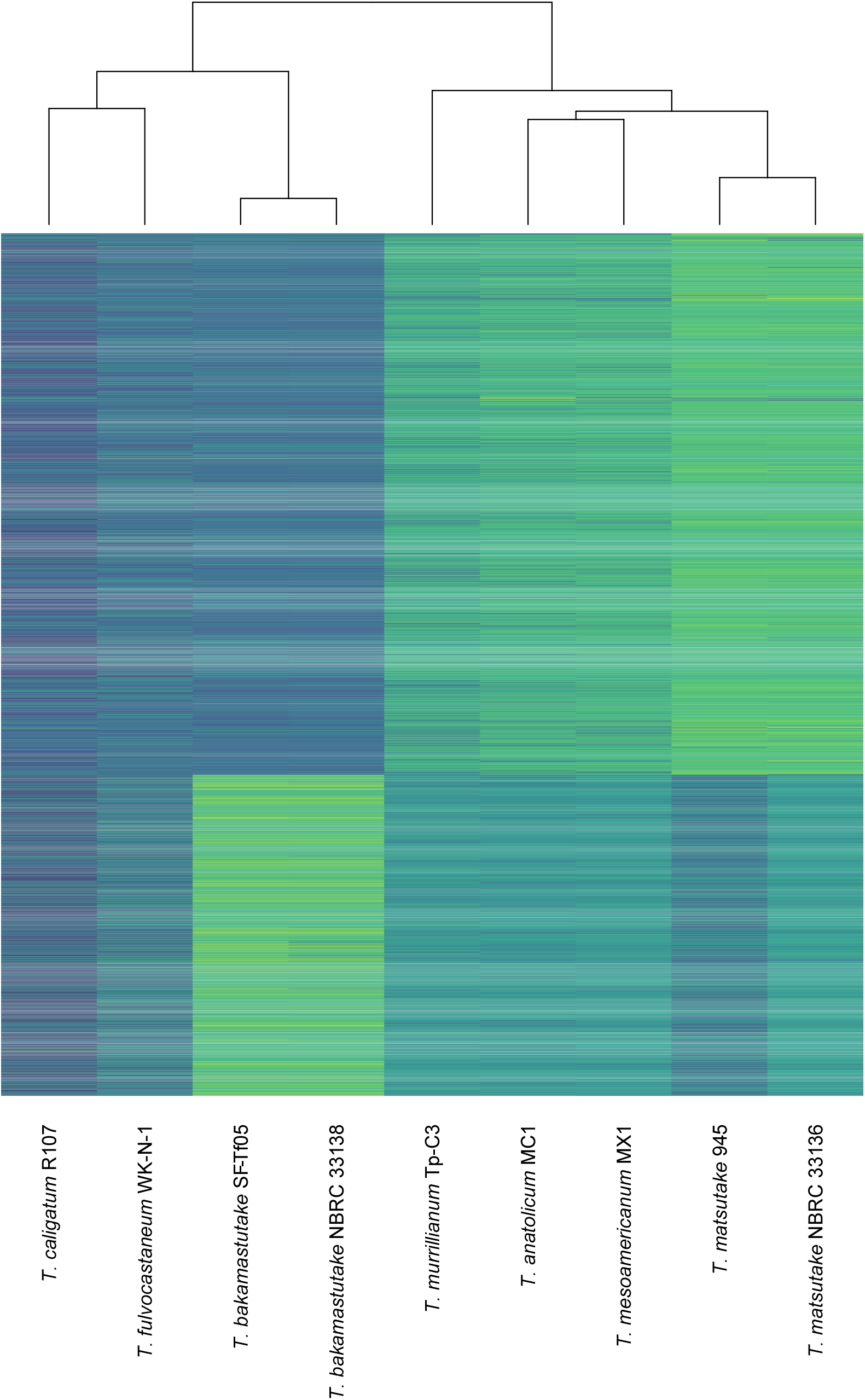
Cluster analysis of *Tricholoma* species according to the presence of shared genes. The default parameters of the heatmap function were used to cluster and visualize the percentages of the covered regions for the predicted genes in Tbkm_v1 and TMA_r1.0. The analysis revealed at least three groups within *Tricholoma* section *Caligata*.

## Discussion

In the present study, we determined the telomere-to-telomere complete genome sequence of the wild-type *T. bakamatsutake* strain SF-Tf05. Similar to the results of our analysis of *T. bakamatsutake* in this study, *T. matsutake* was also assumed to have 13 chromosomes in its nuclear genome. Thus, *Tricholoma* species likely contain 13 chromosomes. Although *T. bakamatsutake* and *T. matsutake* had the same number of chromosomes, the lengths of the corresponding chromosomes differed between these two species. These findings suggest the genome nucleotide contents differentially increased among *Tricholoma* spp. during evolution, but the original number of chromosomes was unchanged. The analysis of conserved non-repeated genes uncovered several inter- and intra-chromosomal rearrangements between *T. bakamatsutake* and *T. matsutake*. Chromosomes 1 and 13 in *T. bakamatsutake* had chimeric structures that partially matched structures in chromosomes 10 and 12 in *T. matsutake*. The comparison with the corresponding chromosomes in *T. matsutake* indicated chromosomes 5 and 12 in *T. bakamatsutake* had multiple inversions. Chromosomal rearrangements, most typically inversions, inhibit synapsis during meiosis, resulting in defective gametogenesis and/or gametophytic lethality. For example, the “balancer chromosomes” in *Drosophila melanogaster*, which were described nearly 100 years ago and have since served as essential genetic resources for studies on fruit flies (Miller *et al*., 2019), contain multiple inverted and rearranged segments and lethal markers for the maintenance of recessive lethal mutations during natural selection. In plants, chromosomal inversions are major factors that facilitated the evolution of reproductive isolation between populations (Baack *et al*., 2015). Accordingly, the chimeric structures and multiple inversions in *T. bakamatasutake* and *T. matsutake* chromosomes may reflect the changes during the differentiation between these two species.

In addition to chromosome-scale structural differences, we detected substantial diversity in the gene contexts between the two examined species. Our gene conservation analysis showed that more than 40% of the *T. matsutake* genes are not present in the *T. bakamatsutake* genome, whereas 83.54% of the *T. bakamatsutake* genes are also present in *T. matsutake*. These observations are consistent with the results of earlier research involving transposon analyses (Murata *et al*., 2013b), which indicated that *T. bakamatsutake* may have diversified earlier than many of the other species belonging to *Tricholoma* section *Caligata*, including *T. fulvocastaneum, T. caligatum, T. merrillianum, T. mesoamericanum, T. anatolicum*, and *T. matsutake*. The diversification of *T. matsutake* may have occurred last. A DNA barcode analysis showed that *T. bakamatsutake* is relatively closely related to *T. matsutake* (Murata *et al*., 2013a). Our gene conservation and read mapping analyses indicated that the genomic structures and gene contexts of *T. caligatum* and *T. fulvocastaneum* differ from those of *T. bakamatsutake* and *T. matsutake*. This is in accordance with the findings of a recent phylogenetic analysis conducted on the basis of internal transcribed spacer (ITS) sequences (Herrera *et al*., 2022). Thus, *T. caligatum* and its related species form an independent group in *Tricholoma* section *Caligata*. Previous studies confirmed *T. dulciolens* is morphologically and phylogenetically distinct from *T. caligatum* (Kytövuori, 1988; Murata *et al*., 2013a), and this is also supported by the phylogenetic relationships according to ITS sequences (Herrera *et al*., 2022). Although we did not analyze *T. dulciolens* and *T. ikkae* in the present study, we speculate that these species form an independent clade. Therefore, *Tricholoma* section *Caligata* likely comprises four major karyotypes.

The distribution of the thoroughly characterized *T. matsutake* retrotransposons *marY1* and *marY2N* also varied between *T. bakamatsutake* (Tbkm_v1) and *T. matsutake* (TMA_r1.0). In *T. bakamatsutake, marY1* was one of the most frequent retrotransposons and was dispersed throughout the nuclear genome, but not in the mitochondrial genome. This is consistent with the results of a previous examination of *T. matsutake* (Kurokochi *et al*., 2022), suggesting that it may be a common feature in the genus *Tricholoma* and/or section *Caligata*. In filamentous fungi (e.g., *N. crassa*), centromeric regions consist of degenerated copies of retrotransposons and simple sequence repeats (Smith *et al*., 2012). Furthermore, LINEs are highly enriched near the center of chromosomes (i.e., centromeric regions) in *T. matsutake*, but a similar localization of general LINEs was not observed in *T. bakamatsutake*, with the exception of a few chromosomes. Instead, at least 10 of the 13 *T. bakamatsutake* chromosomes had *marY2N*-rich regions, often near the center, but sometimes at one end or at multiple locations in which *marY2N* contents exceeded 50% of the analyzed interval (5 kb). An experimental approach, such as ChIP-seq with CenH3, may be necessary to further demonstrate the link between *marY2N* and the centromere function, which has not been determined for any *Tricholoma* species.

## Acknowledgments

The authors thank Yusaku Nishimiya (Plant Genome Evolution Research Team, RIKEN) for his technical assistance. The bioinformatics analysis was performed using the HOKUSAI supercomputing system under project number Q22208 (to HI). The development of the mutation analysis pipeline was partly supported by JSPS KAKENHI (grant number 22K05579 to HI).

## Author Contributions

HI, HM, and SH conceived and designed the experiments. AO and AY collected and provided the fungal strains. HI, HM, and SH performed the experiments. HI designed and performed the bioinformatics analysis and analyzed the data. HI, HM, and SH wrote the manuscript. All authors read and approved the final manuscript.

## Data Availability

The assembled chromosomal and mitochondrial genomic sequences were deposited in GenBank with the accession codes CP114857–CP114870. All other relevant data are provided in the manuscript and the supplementary material.

## Competing Interests

The authors have no competing interests to declare.

## Figure Legends

**Figure S1**. *marY1* contents in the 13 *T. bakamatsutake* SF-Tf05 chromosomes. The *marY1* contents among different chromosomes were determined using a window size of 50 kb. The results indicate that *marY1* is distributed throughout the genome.

**Figure S2**. Distribution of *nit-6* and *SF09g00300* in *T. bakamatsutake* and its allied species. The presence and absence of *nit-6* and *SF09g00300*, encoding a nitrite reductase and a nitrate/nitrite transporter, respectively, in (A) *T. bakamatsutake*, (B) *T. matsutake*, and (C) other *Tricholoma* section *Caligata* species were determined on the basis of a PCR analysis. The geographical locations where the strains were originally collected are indicated in parentheses. The prefectures in Japan are provided in full, whereas three-letter codes are provided for countries (ISO 3166-1 alpha-3).

## Notes

### Competing Interest Statement

The authors have declared no competing interest.

