## Supplementary figures and images for "Complete *de novo* assembly of *Tricholoma bakamatsutake* chromosomes revealed the structural divergence and differentiation of *Tricholoma* genomes"

### Fig. S1

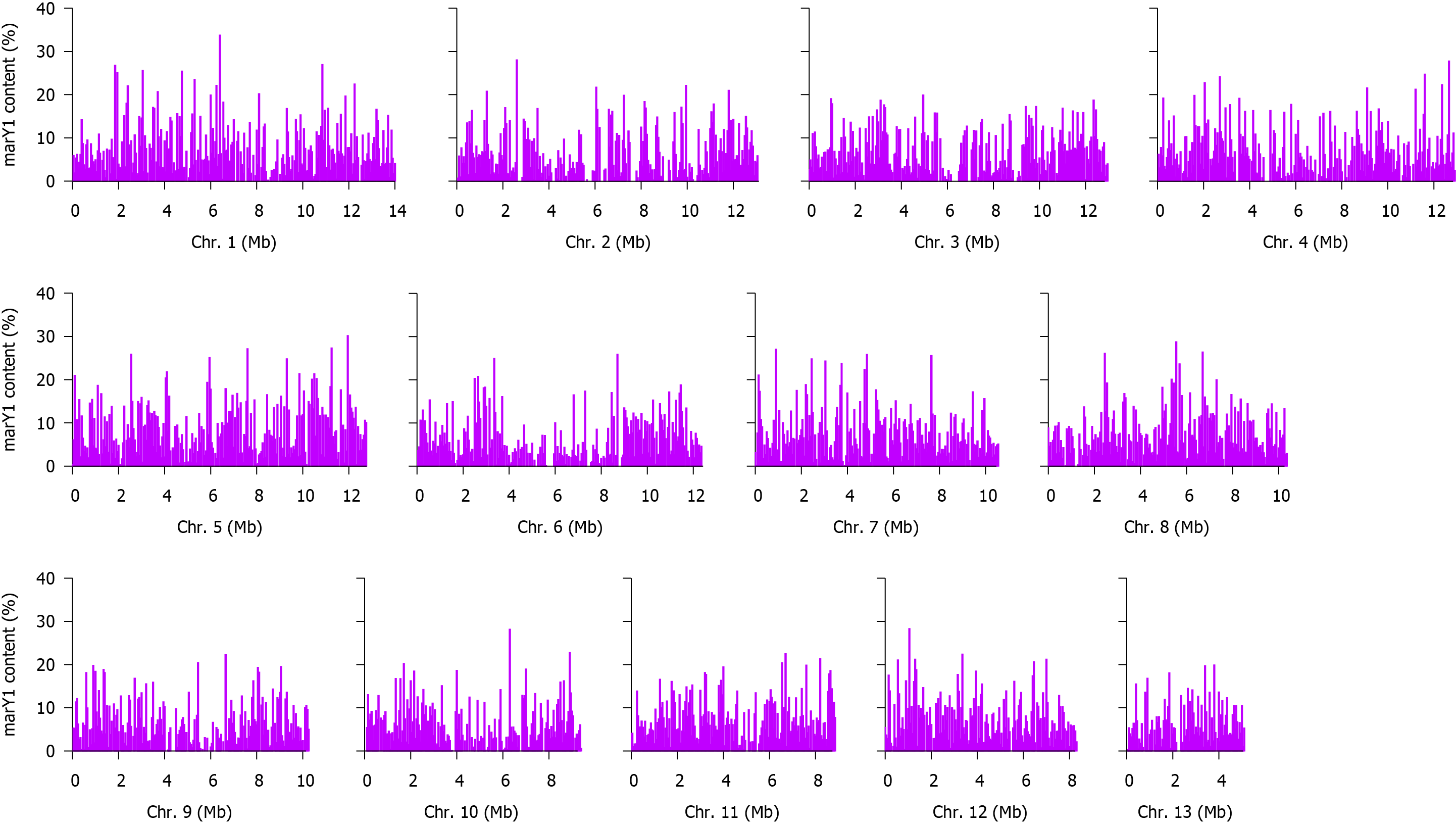

### Fig. S2

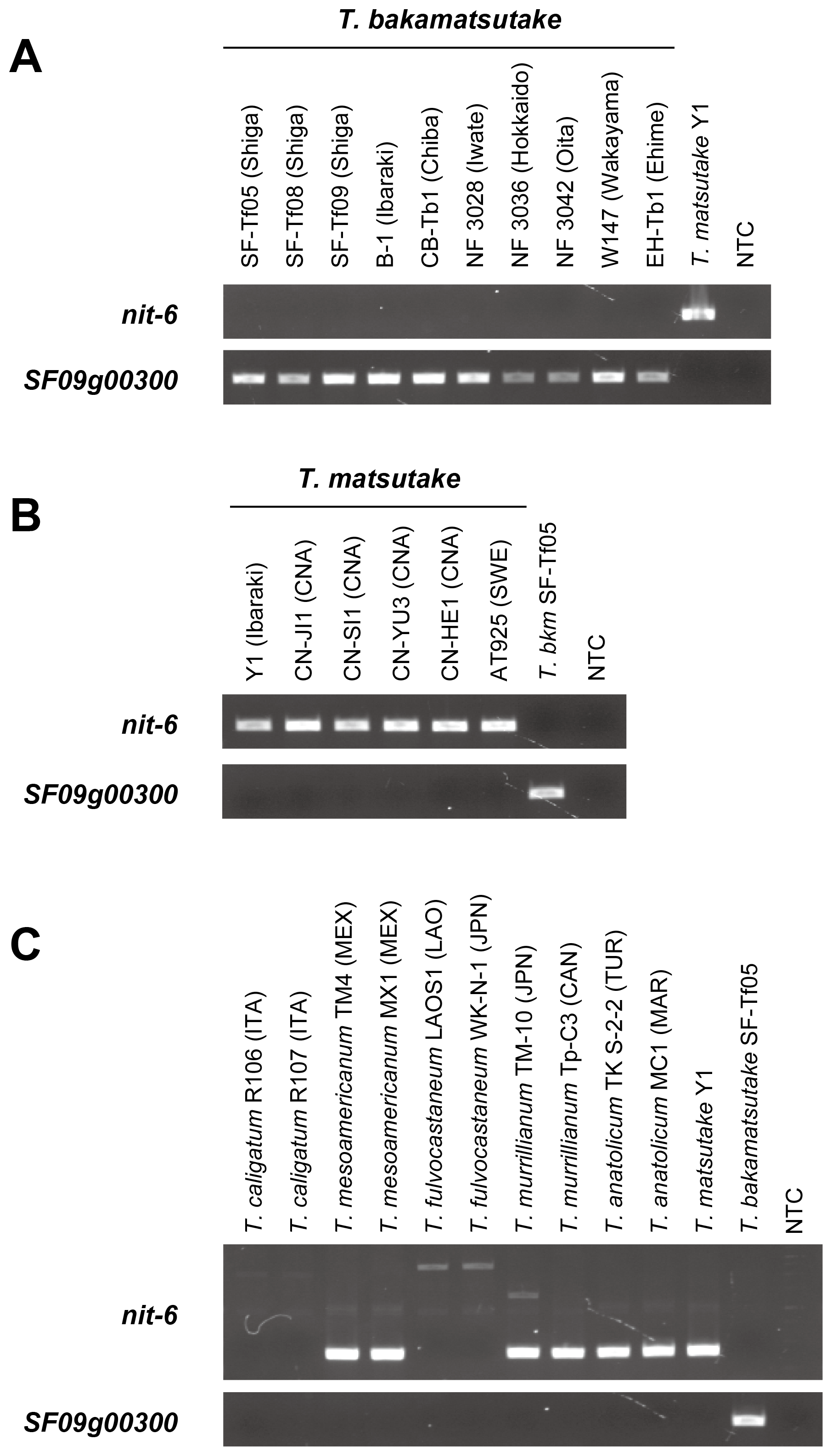
