## Supplementary material for "Complete *de novo* assembly of *Tricholoma bakamatsutake* chromosomes revealed the structural divergence and differentiation of *Tricholoma* genomes": Table S1

Table S1. Genetic statistics for the *T. bakamatsutake* SF-Tf05 chromosomes

| **Chromosome** | Length (bp) | GC (%) | Number of genes | | GenBank accession |
|---|---|---|---|---|---|
| | | | Protein coding | tRNA | |
| **1** | 14,073,764 | 43.79 | 1,071 | 26 | CP114857 |
| **2** | 13,278,086 | 44.52 | 1,342 | 29 | CP114858 |
| **3** | 13,010,463 | 44.10 | 1,125 | 35 | CP114859 |
| **4** | 12,983,161 | 44.34 | 1,203 | 31 | CP114860 |
| **5** | 12,957,022 | 43.72 | 877 | 65 | CP114861 |
| **6** | 12,427,164 | 44.34 | 1,104 | 34 | CP114862 |
| **7** | 10,612,962 | 43.39 | 636 | 30 | CP114863 |
| **8** | 10,437,183 | 43.41 | 668 | 20 | CP114864 |
| **9** | 10,325,240 | 44.13 | 758 | 36 | CP114865 |
| **10** | 9,419,585 | 44.14 | 783 | 13 | CP114866 |
| **11** | 8,909,905 | 43.68 | 600 | 16 | CP114867 |
| **12** | 8,408,117 | 43.45 | 517 | 25 | CP114868 |
| **13** | 5,225,559 | 43.70 | 376 | 12 | CP114869 |
| **Total** | **142,068,211** | **43.94** | **11,060** | **372** | — |
