## Supplementary material for "Complete *de novo* assembly of *Tricholoma bakamatsutake* chromosomes revealed the structural divergence and differentiation of *Tricholoma* genomes": Table S2

Table S2. Categorization of the predicted genes in Tbkm_v1 according to their functions

| KEGG Orthology (KO) entry and functional category | | No. of genes |
|---|---|---|
| ko04131 | Membrane trafficking | 411 |
| ko02000 | Transporters | 209 |
| ko03019 | Messenger RNA biogenesis | 204 |
| ko03036 | Chromosome and associated proteins | 197 |
| ko03009 | Ribosome biogenesis | 180 |
| ko03021 | Transcription machinery | 152 |
| ko01002 | Peptidases and inhibitors | 149 |
| ko04121 | Ubiquitin system | 136 |
| ko03041 | Spliceosome | 131 |
| ko03011 | Ribosome | 128 |
| ko03029 | Mitochondrial biogenesis | 127 |
| ko01001 | Protein kinases | 119 |
| ko03400 | DNA repair and recombination proteins | 96 |
| ko01009 | Protein phosphatases and associated proteins | 95 |
| ko03032 | DNA replication proteins | 80 |
| ko03110 | Chaperones and folding catalysts | 78 |
| ko03016 | Transfer RNA biogenesis | 76 |
| ko03000 | Transcription factors | 67 |
| ko00190 | Oxidative phosphorylation | 66 |
| ko03012 | Translation factors | 59 |
| ko04147 | Exosome | 59 |
| ko01007 | Amino acid related enzymes | 53 |
| ko01003 | Glycosyltransferases | 52 |
| ko01004 | Lipid biosynthesis proteins | 31 |
| ko03051 | Proteasome | 31 |
| ko04812 | Cytoskeleton proteins | 31 |
| ko00010 | Glycolysis / Gluconeogenesis | 30 |
