## Supplementary material for "Complete *de novo* assembly of *Tricholoma bakamatsutake* chromosomes revealed the structural divergence and differentiation of *Tricholoma* genomes": Table S3

Table S3. Corresponding chromosomes between *T. bakamatsutake* (Tbkm_v1) and *T. matsutake* (TMA_r1.0)

| *T. bakamatsutake* chromosome (Tbkm_v1) | *T. matsutake* chromosome (TMA_r1.0) |
|---|---|
| 1 | 10 and 12 |
| 2 | 3 |
| 3 | 5 |
| 4 | 6 |
| 5 | 1 * |
| 6 | 4 |
| 7 | 2 |
| 8 | 9 |
| 9 | 13 |
| 10 | 8 |
| 11 | 7 |
| 12 | 11 * |
| 13 | 10 and 12 |

* Probable intra-chromosomal inversions were detected
