## Supplementary material for "Complete *de novo* assembly of *Tricholoma bakamatsutake* chromosomes revealed the structural divergence and differentiation of *Tricholoma* genomes": Table S4

Table S4. Percentage of the reads that were mapped to the *T. bakamatsutake* and *T. matsutake* reference genome sequences

| **Strain and reference genome** | ***T. bakamatsutake* (Tbkm_v1) %** | ***T. matsutake* (TMA_r1.0) %** |
|---|---|---|
| **"Bakamatsutake" group** | | |
| *T. bakamatsutake* SF-Tf05 | **99.21** | 60.02 |
| *T. bakamatsutake* NBRC 33138 | **97.00** | 58.77 |
| **"Matsutake" group** | | |
| *T. matsutake* 945 | 29.70 | **98.57** |
| *T. matsutake* NBRC 33136 | 51.61 | **96.17** |
| *T. anatolicum* MC1 | 56.80 | **95.54** |
| *T. mesoamericanum* MX1 | 51.26 | **96.19** |
| *T. murrillianum* Tp-C3 | 54.41 | **93.37** |
| **"Caligatum" group** | | |
| *T. caligatum* R107 | 57.22 | 66.28 |
| *T. fulvocastaneum* WK-N-1 | 40.55 | 48.32 |
